# Conformational Reorganization of a Processive Cellulase on Crystalline Cellulose

**DOI:** 10.64898/2026.10.21.760844

**Authors:** Sai Venkatesh Pingali, Junhong He, Loukas Petridis, Barbara R. Evans, William T. Heller, Emal Alekozai, Xiaolin Cheng, Benjamin Lindner, Jeremy C. Smith, Volker S. Urban, Brian H. Davison, Paul Langan, Hugh O’Neill

## Abstract

Enzymes that degrade insoluble polymers must couple catalytic chemistry to productive engagement with heterogeneous substrate interfaces, yet direct structural information on these interfacial states remains limited. We investigated the conformational organization of the processive cellulase *Trichoderma reesei* Cel7A both free in solution and bound to an insoluble cellulose substrate using small-angle neutron scattering (SANS). Use of biosynthetically deuterated bacterial cellulose allowed the scattering contribution of the cellulose-associated Cel7A holoenzyme to be selectively highlighted using SANS with contrast matching. SANS measurements performed under conditions supporting high or low catalytic activity revealed distinct differences between the free and cellulose-bound enzyme. Under poorly active conditions, the conformational distributions of free and bound Cel7A were similar. In contrast, catalytically favorable conditions produced an additional interdomain correlation in the bound enzyme, consistent with redistribution toward a substrate-selected conformational ensemble. Molecular dynamics simulations showed that free Cel7A samples can assume compact, intermediate, and extended conformations, but these solution-state ensembles did not reproduce the structural features of cellulose-bound Cel7A. These results identify productive cellulose engagement as a structurally distinct state of the Cel7A holoenzyme and support a dynamic model in which the linker and carbohydrate-binding module modulate interfacial positioning of the catalytic domain rather than functioning as a rigid mechanical unit during processive hydrolysis. More broadly, the findings suggest that heterogeneous biocatalysis depends not only on substrate binding and active-site chemistry, but also on interface-selected conformational states that determine whether binding becomes productive. The study further demonstrates how deuteration and neutron contrast variation can reveal functional macromolecular conformations directly within complex interfacial environments.

## Introduction

Enzymes that degrade insoluble polymers operate under fundamentally different constraints from enzymes acting on soluble substrates. In addition to catalyzing bond cleavage, these enzymes must first interact with a heterogeneous solid–liquid interface, identify accessible polymer segments or chain ends, and form a catalytically competent complex. Depending on the system, efficient turnover may then require processive movement along the polymer, repeated cycles of dissociation and rebinding, or sustained surface association. Consequently, the overall efficiency of polymer degradation reflects not only active-site chemistry but also substrate accessibility, productive and nonproductive adsorption, polymer-chain engagement, processive motion, and eventual dissociation ^1,2^. These features are evident across enzymes acting on crystalline polysaccharides such as cellulose, chitin, and starch, as well as in emerging studies of enzymes that depolymerize synthetic polymers such as polyethylene terephthalate (PET) ^2–7^.

Bacteria and fungi secrete a variety of enzymes that act synergistically to degrade the crystalline fibrous cellulosic component of plant cell walls into soluble sugars. One of the most extensively studied cellulases is the family 7 cellobiohydrolase Cel7A from *Trichoderma reesei*, which represents a major component of the extracellular cellulase system of this fungus ^8^. Cel7A is a processive enzyme that acts predominantly on free reducing ends of individual cellulose chains exposed at the surface of cellulose fibrils and cleaves cellobiose units as it proceeds ^9,10^. It consists of an N-terminal glycoside hydrolase family 7 catalytic domain (CD) and a C-terminal family 1 carbohydrate-binding module (CBM1), connected by a short, flexible, O-glycosylated, proline/serine/threonine-rich linker. High-resolution structures of the catalytic domain and CBM have been determined independently by crystallography and NMR, respectively, and identified an approximately 50Å long tunnel in the catalytic domain capable of accommodating a single cellulose chain ^11–13^. Direct high-speed atomic force microscopy subsequently demonstrated unidirectional processive movement of Cel7A on crystalline cellulose at approximately 3.5 nm s⁻¹. Notably, the isolated catalytic domain moved at a similar velocity, indicating that the catalytic domain itself contains the essential machinery for processive translocation, whereas the CBM and linker modulate interactions of the holoenzyme with the cellulose interface ^14^.

Despite intensive study, the binding mechanism of *T. reesei* Cel7A to cellulose by both the CD and CDM is not understood fully and cannot be described simply as formation of a single enzyme– substrate complex. Recent single-molecule measurements have shown that cellulose-bound Cel7A molecules populate kinetically distinct states. Haviland et al. tracked more than 11,000 individual quantum dot labeled Cel7A molecules and found that most surface-associated molecules remained static during the observation period, whereas a smaller population underwent processive motion; individual enzymes could also transition between static and moving states ^15^. These observations support a mechanistic distinction between adsorption to the cellulose surface, productive complexation of a cellulose chain within the catalytic tunnel, processive hydrolysis, and stalled or nonproductive binding. The individual steps can also be perturbed independently. For example, single-molecule analysis of product inhibition showed that cellobiose both decreases the initial association of Cel7A with cellulose and reduces the velocity and run length of enzymes that have already entered a processive state ^16^. Thus, an important unresolved question is how productive engagement with cellulose is reflected in the structure and interdomain organization of the intact Cel7A holoenzyme.

The flexible linker and CBM are likely to be important in defining this interfacial conformational landscape. Early models proposed that the linker coordinates the catalytic domain and CBM through compression–extension motions that allow the two domains to advance along cellulose in a “caterpillar-like” manner, with the CBM helping to localize or extract cellulose chains while the catalytic domain performs processive hydrolysis ^17–19^. Deletion or removal of the linker and CBM can reduce binding and activity on crystalline cellulose, demonstrating that these regions make important contributions to holoenzyme function ^20^. However, the observation that the isolated catalytic domain is itself capable of processive motion argues against a requirement for the CBM to act as a mechanical motor during every catalytic step ^14^. Instead, subsequent computational and experimental work has emphasized the dynamic nature of the multidomain enzyme. Simulations of full-length Cel7A indicate that O-glycosylation of the linker restricts its conformational ensemble, favors more extended conformations, and promotes coordinated motion between the catalytic domain and CBM ^21^. These findings suggest that the linker and CBM may regulate the probability and geometry of productive interfacial engagement rather than functioning simply as a mechanical spring. Although atomic level crystallographic information on the holoenzyme has so far been elusive, its solution conformation has been studied with SAXS and SANS revealing insights into pH dependent conformational changes that can be correlated to its catalytic activity and the effect of O-glycosylation on the linker domain on the interaction of the CD and CBM ^22–24^. Nevertheless, direct experimental structural information on the intact Cel7A holoenzyme while associated with insoluble cellulose remains limited.

A major technical challenge in investigating enzyme action at heterogeneous biological interfaces is the difficulty in obtaining molecular-level information when the enzyme is bound to an insoluble substrate. Small-angle neutron scattering (SANS) provides a means of overcoming this limitation because isotopic substitution can generate contrast between enzyme and the insoluble substrates. This was demonstrated in a recent study that investigated a deuterium-labeled multidomain lytic polysaccharide monooxygenase binding to a chitin substrate ^25^. In the present study, bacterial cellulose was biosynthetically deuterated ^26,27^ so that its scattering contribution could be strongly attenuated by contrast matching, allowing the scattering from cellulose-associated *T. reesei* Cel7A to be examined independently. We used contrast-variation SANS together with molecular dynamics simulations to compare the conformational ensembles of Cel7A free in solution and bound to cellulose under conditions that support either high or low catalytic activity. Specifically, we asked whether cellulose association alters the interdomain organization of the intact enzyme, whether catalytically active and poorly active binding conditions produce distinguishable conformational ensembles, and whether the resulting structures are consistent with models in which the catalytic domain and CBM remain coordinately engaged with the same cellulose chain during processive hydrolysis.

## Materials and Methods

### Production and purification of cellulase and cellulose

Propagation of *Acetobacter xylinus sub sp. sucrofermentans* (ATCC 700178) in D_2_O-based media and purification of deuterated bacterial cellulose has been described ^26–28^. The final growth medium contained ∼97.6% D_2_O media with *H*_8_-glycerol as the carbon source and produced 0.5 g dry cellulose / L of culture. This growth condition yielded a contrast match point of 85.4 ± 1.5 % D_2_O for the deuterated cellulose (Figure S1 and ^29^).

### Cellulose – Cel7A sample preparation for SANS experiments

*T. reesei* Cel7A was purified from a commercial culture filtrate produced by fermentation of *T. reesei* QM9414 (Sigma-Aldrich Chemical Company, St. Louis, MO, USA) as described previously ^24,30–32^. The experiments describing the determination of optimal Cel7A binding to bacterial cellulose are described in the Supplementary Information (Table S1, Figures S2 and S3). For preparation of the SANS samples, the cellulose hydrogels (∼80g) were frozen, placed in 20 mL of H_2_O, and disrupted using a blender (Osterizer Mo 83-1) until a homogeneous slurry was formed (approx. 2 min). The ground cellulose was collected by centrifugation (4500 x g for 20 min) and resuspended in H_2_O. Initially, the cellulose was incubated and exchanged in 0.1N NaOD in D_2_O three times to exchange the labile hydrogens for deuterium. This step was followed by three sequential washings with 50 mM Na acetate buffer at pD (pD = pH + 0.4) values of 4.2, 5.3, or 7.0 containing the appropriate ratio of D_2_O/H_2_O at ∼8 to 10 h intervals to equilibrate the cellulose and neutralize the NaOD. For SANS measurements, the samples were prepared by mixing ∼0.2 mL of 50 mg/mL cellulose with 0.6 mL of ∼5 mg/mL Cel7A and incubated in a water bath for 15 min at 15 °C. This was followed by 3 sequential centrifugation (16000 x g for 60 sec) and resuspension steps to remove the unbound enzyme. After the final wash, the sample was resuspended in the buffer and transferred to a 1mm path cylindrical Hellma quartz cuvette (Model# 120-QS 1.0 mm) for measurement. The supernatant solutions that were recovered during washing were analyzed for protein and sugar content as described above. For samples containing ∼10 – 11 mg cellulose, approximately 35.5, 51.5 and 55.3% of the initial Cel7A added were bound to the cellulose at pD 4.2, 5.3, and 7.0, respectively.

### Small-angle neutron scattering

SANS measurements were performed with the CG-3 Bio-SANS instrument ^33^ at the High Flux Isotope Reactor (HFIR) facility of Oak Ridge National Laboratory. The three different instrument configurations employed to cover the range, 0.003 < *Q* (Å^−1^) < 0.4, of scattering vectors using sample-to-detector distances of 1129 mm, 6829 mm and 15329 mm. The detector (1 m x 1 m GE-Reuter Stokes Tube Detector) was offset by 350 mm from the beam. *Q* = (4*π*/*λ*)sin*θ* is the scattering vector, 2*θ* is the scattering angle and λ is 6 Å (*λ*⁄Δ*λ* = 0.132).

The scattering intensity profiles *I*(*Q*) versus *Q*, were obtained by azimuthally averaging the processed 2D images, which were normalized to incident beam monitor counts, and corrected for detector dark current, pixel sensitivity and scattering from backgrounds: D_2_O buffer and quartz cell.

The SANS measurements were carried out at 15 °C to minimize hydrolysis of the cellulose while ensuring that sufficient amount of enzyme remains bound to the cellulose for the duration of the measurement, a compromise that has been recommended for investigation of binding by cellulases ^34^. The scattering intensity profiles *I*(*Q*) versus *Q*, for Cel7A bound to bacterial cellulose were obtained by taking 5 min exposures over a span of 20 min. All 5 min exposures resulted in the same profile and were summed into one 20 min exposure for reduction and analysis. The radius of gyration, *R_g_*, was determined according to Guinier and Fournet ^35^. Because subtraction of the residual cellulose scattering increased uncertainty in the lowest-Q region in the data for cellulose-bound Cel7A, Guinier-derived parameters which resulted in large error bars were not used for quantitative analysis of cellulose-bound Cel7A. Structural comparisons of the bound enzyme were instead based on indirect Fourier transformation of the scattering profiles implemented in GNOM ^36^, which incorporates information over a broader Q-range and provides the real-space distance distributions (*R_g_* and maximum dimension (*D_max_*). This approach has been previously recommended for interpretation of SANS data of multicomponent systems ^37^. Structural modeling of Cel7A from the SANS data was accomplished using DAMMIN ^38^. A set of 37 models was generated using default settings from each SANS dataset and a consensus envelope, representing the structure consistent across the set of models was determined using DAMAVER program suite ^39^. The Supplementary Information section provides further details on data collection and analysis.

### Molecular dynamics simulations

MD simulations were performed using the CHARMM force field for proteins ^40–42^, and carbohydrates ^43,44^ and the TIP3P model for water ^45^. Theoretical SANS profiles were calculated employing the SASSENA software ^46^. The SI contains details on the model and simulation and how we obtained populations of Cel7A via a procedure seeking to optimize the theoretical scattering profiles against experimental SANS data ^47^. The total theoretical profile, *I_T_*, is the sum of *I*(*Q*) of each cluster, weighted by its population 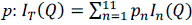 (40). The *p_n_* values of each cluster that together are consistent with the experimental SANS data were then determined by employing an optimization algorithm that minimized the function: 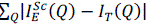.

## Results

### SANS analysis of Cel7A in solution and bound to cellulose

Our overall aim was to use SANS to gain insight into the relative organization of the catalytic and cellulose-binding domains of Cel7A when the enzyme is associated with a cellulose substrate. SANS can distinguish the scattering contributions of individual components in a complex system provided that sufficient contrast exists between them. However, the neutron scattering length densities of *T. reesei* Cel7A and cellulose are similar, equivalent to approximately 40% D₂O solvent, which precludes selective observation of the enzyme using natural-abundance isotopic materials. Bacterial cellulose produced by *Gluconoacetobacter* sp. ^48–51^, a genus later reclassified as *Komagataeibacter* ^52^. was chosen for these experiments because it has a cellulose I crystalline form resembling that of plant cellulose ^50^ and can be produced in highly deuterium-labeled form while maintaining fiber morphology, molecular weight, and crystallinity similar to protiated control material ^26–28^. The scattering from partially deuterated bacterial cellulose was strongly attenuated in D₂O-containing buffer, allowing the scattering contribution of cellulose-associated Cel7A to be measured with minimal contribution from the cellulose substrate.

SANS experiments were performed at three pD values to compare the structure of Cel7A free in solution and bound to cellulose under conditions spanning high and low catalytic activity. SANS profiles of free *T. reesei* Cel7A recorded over the Q-range 0.008 < Q < 0.35 Å⁻¹ at pD 4.2, 5.3, and 7.0 are shown in Figure 1A. Guinier analysis of the low-Q region indicated that the protein was monodisperse and free of detectable aggregation at all three pD values (Figure 2A). The radius of gyration (*R_g_*) and maximum dimension (*D_max_*) obtained from the pair-distance distribution function (*p(R)*) analysis using GNOM ^36^ are summarized in Table 1.

**Figure 1.**
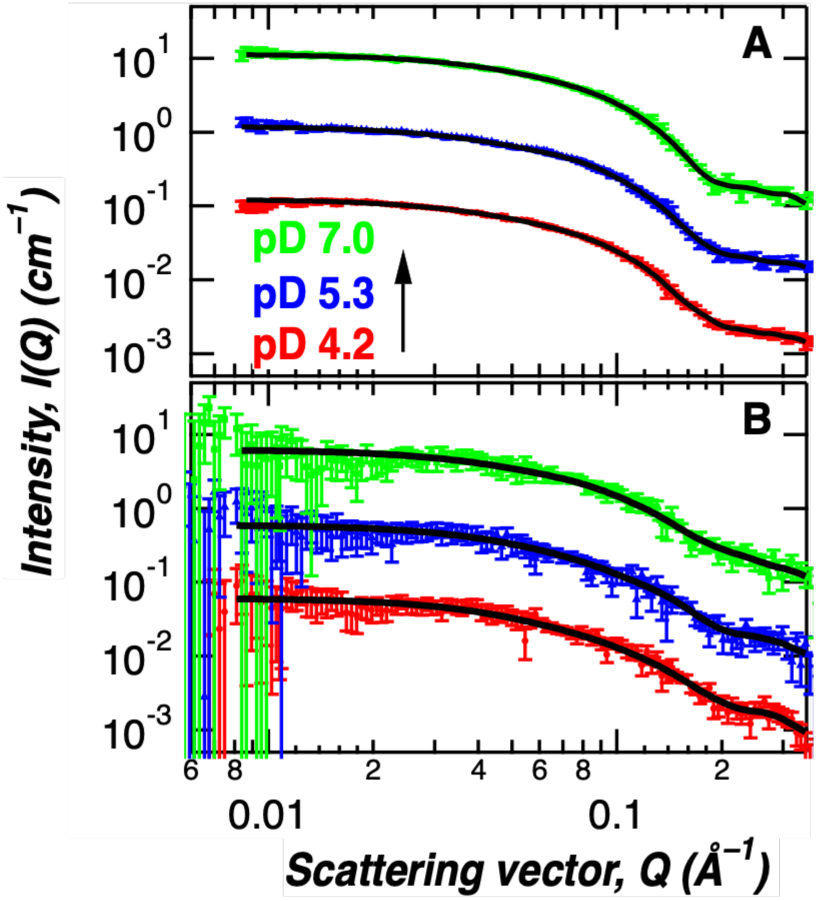
SANS profiles of Cel7A in solution and bound to cellulose. The enzyme was in sodium acetate buffers at pD = 4.2, 5.3, and 7.0 in (A) free and (B) bound states. Curves represent pD= 7.0 (green squares), 5.3 (blue triangle), 4.2 (red dots) and GNOM fit (solid black line)

**Figure 2.**
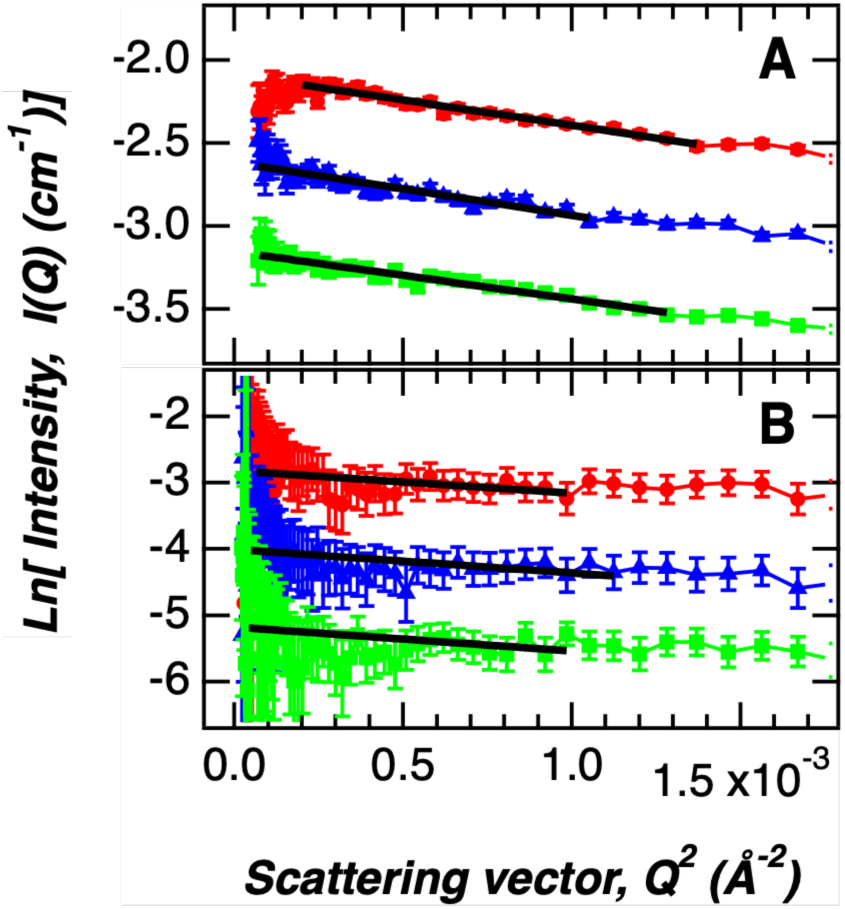
Guinier plots of (A) free and (B) bound Cel7A enzyme SANS curves plotted as a function of pD. Curves represent pD = 7.0 (green squares), 5.3 (blue triangle), 4.2 (red dots) and the *R_g_* values obtained from the slopes (solid black lines) are summarized in Table S4.

**Table 1.** Structural parameters of *T. reesei* Cel7A in solution and bound to cellulose.

| $p(R)$ (Å) | | | | |
| --- | --- | --- | --- | --- |
| pD | $R_g$ | $\pm\Delta R_g$ | $D_{max}$ | $\pm\Delta D_{max}$ |
| 1CEL | 16.51 | 0.04 | 70 | 5 |
| *Free* |  |  |  |  |
| 4.2 | 32.0 | 0.8 | 135 | 5 |
| 5.3 | 32.6 | 0.7 | 130 | 5 |
| 7.0 | 29.4 | 0.6 | 115 | 5 |
| *Bound* |  |  |  |  |
| 4.2 | 30.5 | 1.3 | 115 | 5 |
| 5.3 | 30.2 | 2.2 | 120 | 5 |
| 7.0 | 31.3 | 1.7 | 130 | 5 |

For free Cel7A, all three *p(R)* distributions contain a pronounced peak centered at approximately 24 Å followed by a broad distribution extending to longer distances (Figure 3). The position and shape of the *p(R)* profiles at shorter correlation distances (<50 Å) agree closely with the *p(R)* distribution calculated from the crystal structure of the Cel7A catalytic domain (PDB 1CEL) ^12^ using CRYSON ^53^, indicating that this portion of the distribution is dominated by intradomain correlations within the catalytic domain.

**Figure 3.**
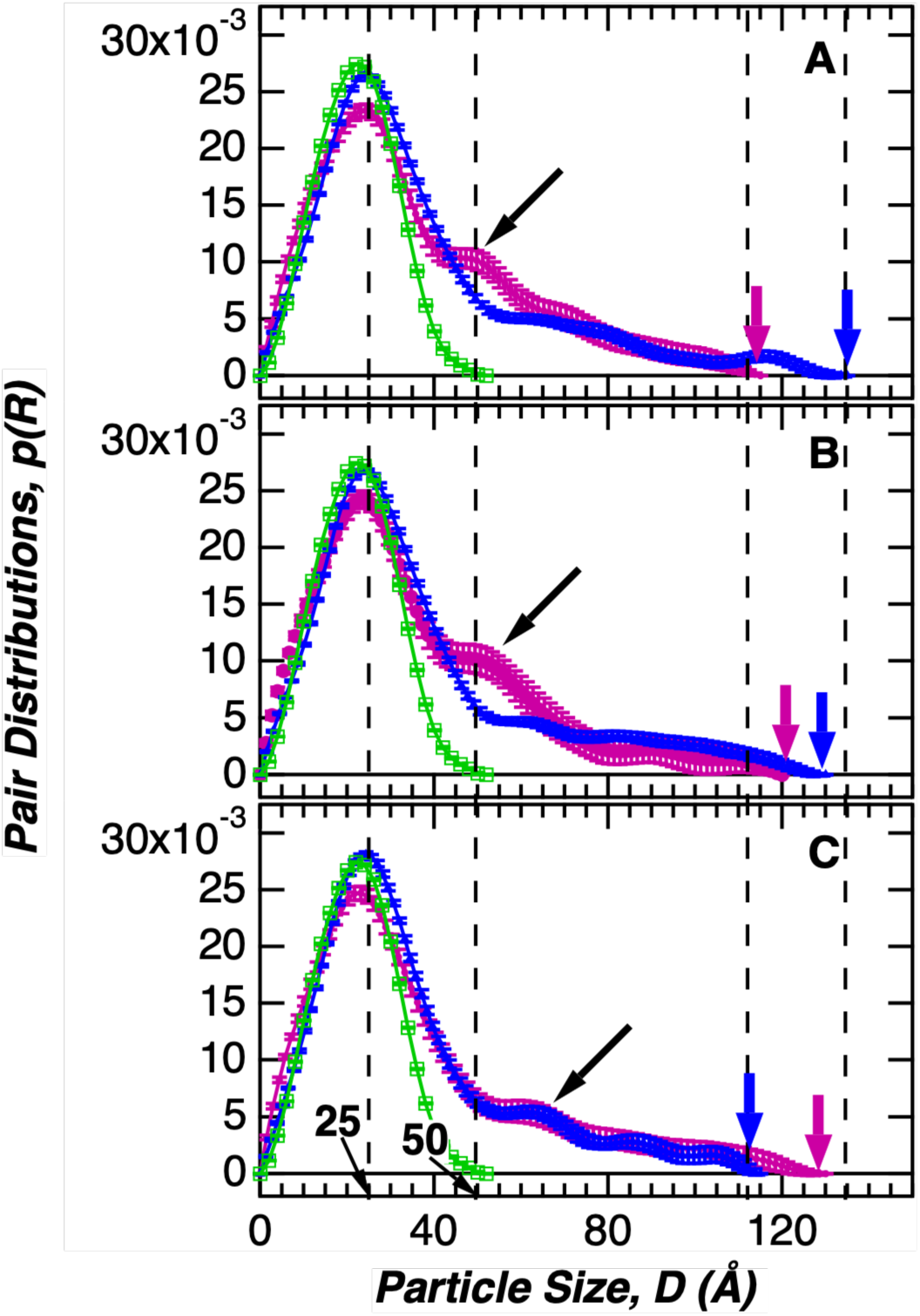
Pair distribution function, *p*(*R*) for Cel7A bound and free in solution. Cel7A was in sodium acetate buffer pD (A) 4.2; (B) 5.3; and (C) 7.0. The green open squares and solid line represents the *p*(*R*) function of only the catalytic core obtained using the published crystal structure 1CEL. The *p*(*R*) for bound Cel7A is shown as purple dots and solid line and the *p*(*R*) for the free protein is shown as blue triangles and solid line. The blue and purple arrows between 110 Å and 135 Å provide a visual guidance in the *D_max_* trends of free and bound Cel7A enzyme respectively, as a function of pD values and summarized in Table 1.

The longer-distance portion of the *p(R)* distributions therefore contain contributions from correlations involving the linker and CBM relative to the catalytic domain. A weak secondary feature centered at approximately 65 Å is also apparent in the free-enzyme distributions (Figure 3). The *D_max_* values are consistent with the CD and CBM being predominantly extended along the axial length of the molecule. The difference in the *D_max_* of the holoprotein and the catalytic core is 65, 60, and 45 Å at pD values 4.2, 5.3 and 7.0, which represents the contribution of the 63 additional amino acids of the linker and CBM relative to the 434 residues of the catalytic core (**Table 1**). These dimensions are substantially shorter than would be expected for a fully extended linker, which has a maximum calculated length of approximately 92 Å ^54^, indicating that the holoenzyme is not fully extended in solution. The *R_g_* and *D_max_* values also demonstrate a pD-dependent change in the average dimensions of free Cel7A, with the largest *D_max_* observed at pD 4.2 and the smallest at pD 7.0.

For cellulose-bound Cel7A, residual low-Q scattering from the contrast-matched cellulose was corrected as described in the Methods and Supplementary Information (**Figure S4**). The resulting scattering profiles are shown in **Figure 1B**, with corresponding Guinier plots in **Figure 2B** and *p(R)* distributions in **Figure 3**. Because this subtraction increased uncertainty in the low-Q region resulting in low confidence in Guinier-derived *R_g_* values, structural comparisons of the bound enzyme were based primarily on the *p(R)* distributions obtained from indirect Fourier transformation. The bound enzyme retained the principal *p(R)* peak at approximately 24 Å under all three conditions, but the longer-distance portion of the distribution showed pronounced pD-dependent differences. The *D_max_* values of cellulose-bound Cel7A were 115, 120, and 130 Å at pD 4.2, 5.3, and 7.0, respectively (**Table 1**), displaying the opposite pD-dependent trend to that observed for free Cel7A. At pD 7.0, the *p(R)* distributions of free and bound Cel7A were similar within the resolution of the experiment, including both the principal ∼24 Å feature and the longer-distance portion of the distributions (**Figure 3C**). In contrast, at pD 4.2 and 5.3, a distinct feature centered at approximately 46 Å appeared in the *p(R)* distributions of the cellulose-bound enzyme (**Figure 3A,B**). This feature was not apparent in the corresponding free-enzyme distributions or in the pD 7.0 bound-enzyme distribution.

The ∼46 Å feature occurs at distances greater than those dominated by the catalytic domain and therefore reflects correlations involving the catalytic domain, linker, and/or CBM. The separation between the principal ∼24 Å peak and this additional feature is approximately 22–24 Å at pD 4.2 and 5.3. A longer-distance feature at approximately 65 Å also remains evident in the bound-enzyme distributions, demonstrating that the bound samples contain a range of interdomain distances rather than a single discrete configuration. A previous SAXS study of *Bacillus subtilis* cellulase 5A, an enzyme of similar overall size and modular architecture, also identified correlation features near 25 and 65 Å and showed that the intensity of the longer-distance feature was sensitive to changes in linker flexibility ^55^. These observations were attributed to changes in correlations between the catalytic and cellulose-binding domains.

### Low-resolution structural analysis of Cel7A

The SANS data were further analyzed using DAMMIN shape-restoration analysis ^38^ to generate low-resolution dummy-atom models of Cel7A free in solution and bound to cellulose at the different pD values (Figure 4). The structures obtained for free Cel7A are consistent with previous solution-scattering and electron-microscopy studies ^22,23,56^. The models are elongated and contain two spatially separated regions. The dimensions of the larger region are similar to those of the high-resolution structure of the *T. reesei* Cel7A catalytic domain ^12^, which is shown superimposed on the dummy-atom models in Figure 4. The remaining density is attributable to the linker and CBM. The low-resolution envelopes of cellulose-bound Cel7A retained the same overall elongated architecture, but differences were observed as a function of pD (Figure 4). At pD 7.0, the free and cellulose-bound models were similar, with well-defined separation between the catalytic-domain and linker/CBM regions. At lower pD values, the dimensions of the catalytic-domain region remained similar to those of the free enzyme, whereas the density associated with the linker and CBM was less well defined in the bound-enzyme models. These differences are consistent with the changes observed in the corresponding *p(R)* distributions (Figure 3).

**Figure 4.**
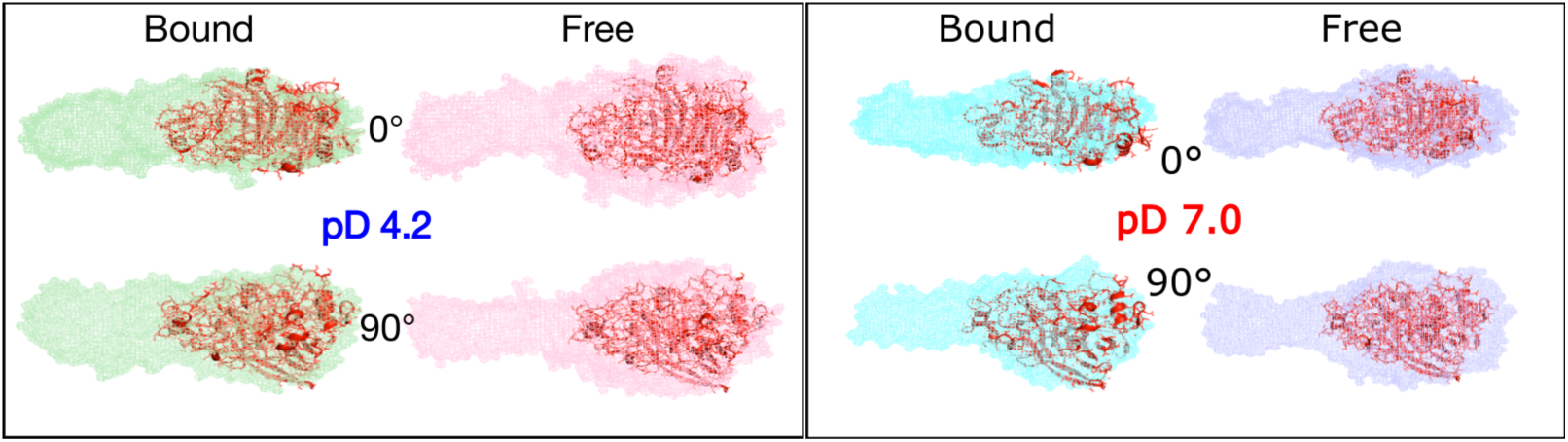
Low resolution structural envelopes obtained using DAMMIN for free and bound states of Cel7A enzyme in its most and least active conditions, pD 4.2 (*left panel*) and 7.0 (*right panel*), respectively.

### Molecular dynamics analysis of Cel7A conformational ensembles

Molecular dynamics simulations were used to characterize the conformational space sampled by full-length Cel7A in solution and to compare the calculated scattering from these conformations with the experimental SANS data. Approximately 20,000 Cel7A configurations generated during the simulations were initially grouped by structural similarity and reduced to 11 representative clusters with distinguishable calculated scattering profiles (Figure S5, S6) ^47^. The theoretical SANS profile of each cluster was calculated and the cluster populations were optimized against the experimental scattering data. For free Cel7A, the experimental SANS profiles at all three pD values could be described using only three of the 11 representative clusters (Figure 5A–D). These three clusters represent relatively compact, intermediate, and extended conformations. The same three structural classes were sufficient to describe the data at each pD, although their relative populations differed (Table 2).

**Figure 5.**
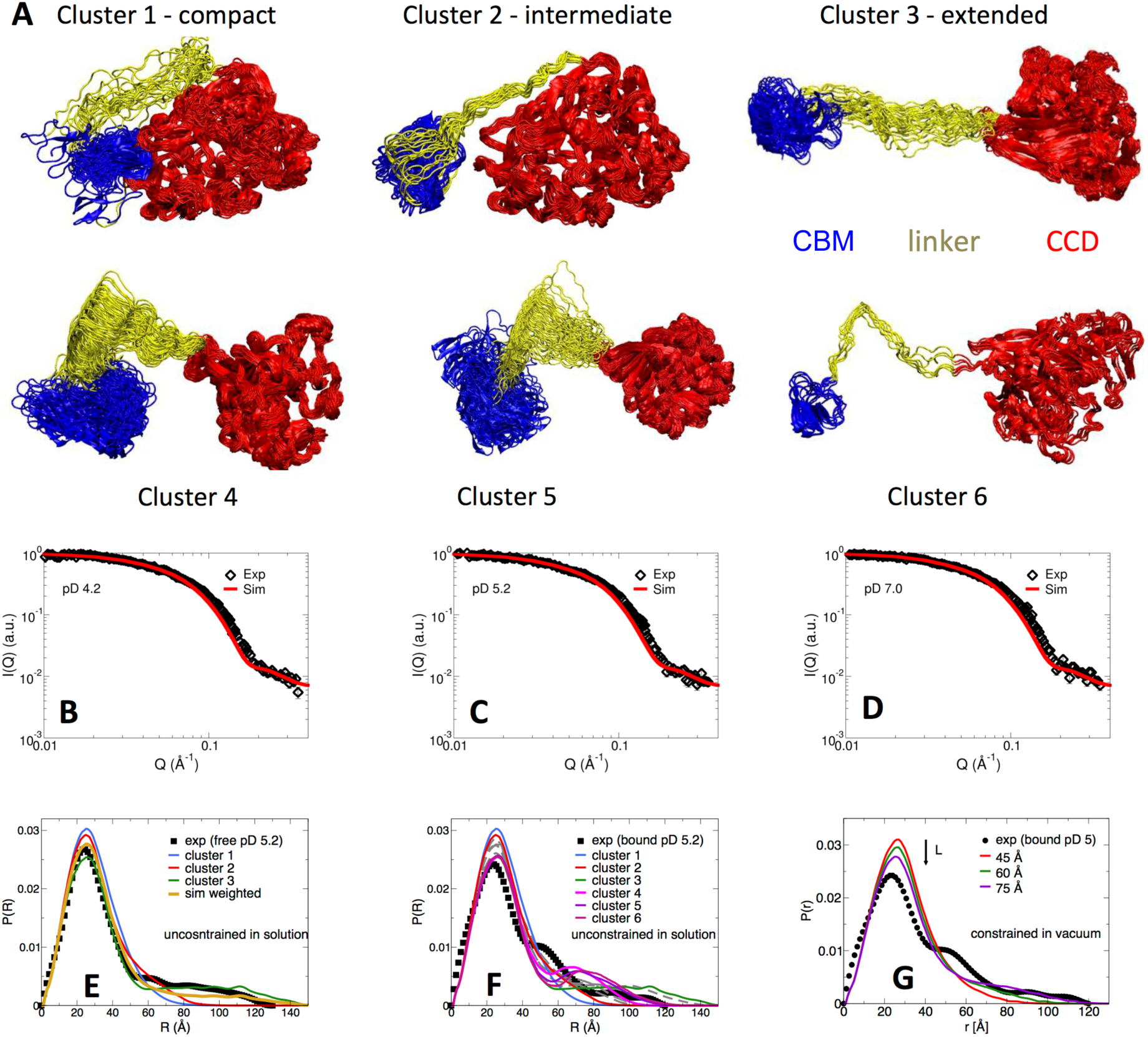
MD simulations of Cel7A free in solution. (A) Clusters sampled by the MD simulation of Cel7A free in solution. Only the top three have non-zero equilibrium populations, when comparing to the SANS experiments. (B)–(D) Comparison of experimental and theoretical (weighted by the populations of Table 2) SANS profiles of free Cel7A at pD of 4.2, 5.3 and 7.0. (E)–(F) *p*(*R*) curves of the six clusters above. (G) *p*(*R*) curves from simulations where the CD–CBM distance is constrained to *L*=45, 60, and 75 Å.

**Table 2.** Equilibrium populations of the MD simulation-derived clusters of Cel7A free in solution as determined by fitting SANS data.

| pD | Cel7A populations (%) |  |  |
| --- | --- | --- | --- |
|  | Compact | Intermediate | Extended |
| 4.2 | 12 | 44 | 44 |
| 5.3 | 36 | 13 | 51 |
| 7.0 | 28 | 29 | 43 |

At pD 4.2, the optimized populations of the compact, intermediate, and extended clusters were 12%, 44%, and 44%, respectively; at pD 5.3 they were 36%, 13%, and 51%; and at pD 7.0 they were 28%, 29%, and 43%, respectively (**Table 2**). Thus, the SANS/MD analysis indicates that free Cel7A exists as an ensemble of multiple interdomain configurations rather than a single average structure. Differences among the experimental profiles at the three pD values can be reproduced primarily through changes in the relative populations of these conformational states. An equivalent population analysis using the conformations sampled in the solution MD simulations but without adjusting populations sizes to fit experiment did not reproduce the experimental scattering from cellulose-bound Cel7A. Of the 11 solution-derived clusters, clusters 4–6 displayed secondary features in their calculated *p(R*) distributions that qualitatively resembled the additional correlation observed experimentally for bound Cel7A at pD 4.2 and 5.3 (Figure 5E, F). However, these calculated secondary correlations occurred at longer distances, approximately 75–80 Å, relative to the experimentally observed feature near 46–50 Å. A common characteristic of these clusters was a pronounced bend or kink in the linker, resulting in multiple linker residues occupying similar distances from the catalytic domain (Figure S6A). To identify the structural origin of the calculated secondary feature, *p(R)* distributions were generated for cluster 4 after selectively including or excluding correlations among the catalytic domain, linker, and CBM. These calculations showed that the secondary feature was dominated by interdomain correlations between the catalytic domain and linker (Figure S6B,C).

To further examine configurations of cellulose-bound Cel7A, additional simulations were performed in which the catalytic domain and CBM were positioned at a cellulose microfibril surface (Figure S7A). In these simulations, the catalytic cleft of the CD faced the cellulose surface and the CD and CBM were constrained to interact with the same cellulose strand, while the glycosylated linker remained unconstrained. The center-of-mass distances between the CD and CBM were varied from 45 to 70 Å in 5 Å increments (Figure S7). The calculated *p(R)* distributions from all of these constrained models contained a principal peak at approximately 25 Å, comparable to the dominant feature in the experimental data (Figure 5G; Figure 6). However, none of the constrained configurations reproduced the additional correlation near 46–50 Å observed experimentally for cellulose-bound Cel7A at pD 4.2 and 5.3, nor did they generate the longer-distance features present in the experimental distributions. Thus, within the configurations sampled here, models in which the catalytic domain and CBM were simultaneously constrained to the same cellulose strand were unable to reproduce the experimental SANS data for the bound enzyme.

**Figure 6.**
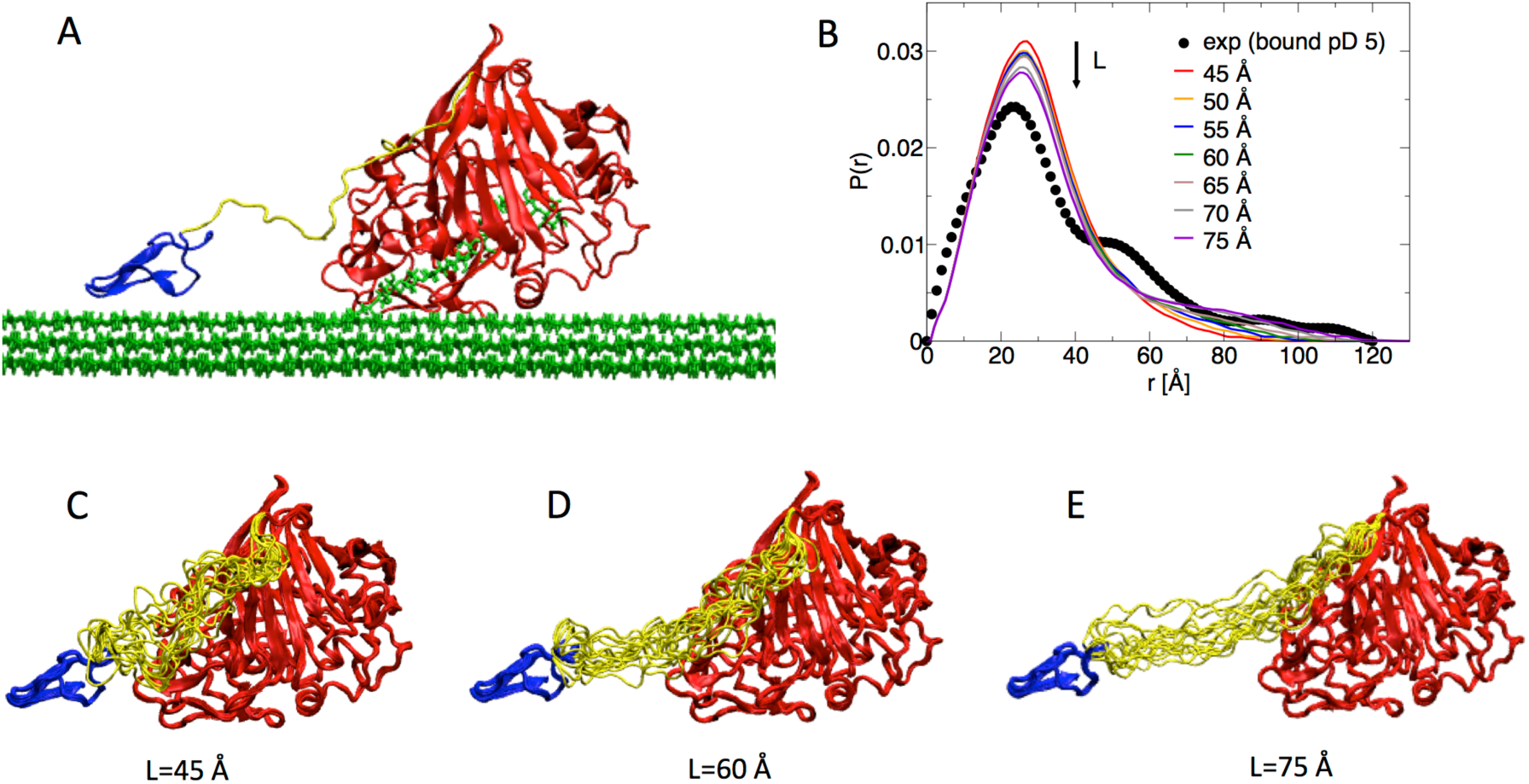
(A) The conformation of Cel7A bound to cellulose used as a template for the constrained simulations of Cel7A in vacuum. The Cα atoms of the catalytic and CBM domains were fixed during these simulations, while the linker and glycosylation residues were unconstrained. Seven simulations were performed, where the center of mass distance between the catalytic and CBM was fixed at *L*=45, 50, 55, 60, 65, 70, and 75 Å. (B) *p*(*R*) curves derived from the seven constrained simulations and (C–E) snapshots of the simulations at *L*=45, 60, and 74 Å are shown in C, D and E, respectively.

## Discussion

In this study, we used contrast-variation SANS in combination with molecular dynamics simulations to investigate the conformational organization of *T. reesei* Cel7A both free in solution and associated with an insoluble cellulose substrate. The ability to selectively attenuate the scattering contribution from deuterated bacterial cellulose provides structural information on the intact holoenzyme while it is bound to cellulose, a state that remains difficult to access using conventional high-resolution structural methods. Analysis of the SANS curves shows significant differences between free and bound Cel7A that provide insight into the relative organization of the catalytic domain (CD), linker, and carbohydrate-binding module (CBM) during cellulose hydrolysis. In particular, the conformational distributions of free and cellulose-bound Cel7A are similar at pD 7.0, where the enzyme binds cellulose but has low catalytic activity, whereas marked differences emerge at pD 4.2 and 5.3, conditions that support cellulose hydrolysis. Under these catalytically favorable conditions, the bound enzyme exhibits an enhanced correlation at approximately 46 Å that is not apparent for either free Cel7A or cellulose-bound Cel7A at pD 7.0. Together with the persistence of longer-distance correlations and similar overall maximum dimensions, these observations suggest that cellulose-bound Cel7A does not adopt a single compact structure but instead populates an ensemble containing both compact and more extended interdomain configurations.

The distinction between the conformational behavior observed under active and poorly active conditions is particularly relevant in light of recent evidence that cellulose-bound Cel7A exists in multiple kinetically distinct states. Single-molecule fluorescence measurements have shown that the majority of surface-associated Cel7A molecules remain static over the period of observation, whereas a smaller population undergoes processive motion; individual enzymes can also reversibly transition between static and processive states before dissociation ^15,57^. Haviland et al. therefore proposed that the transition from a static surface-bound state into a processive state represents an important kinetic limitation in cellulose degradation. More recently, single-molecule measurements of product inhibition demonstrated that cellobiose can inhibit both the initial interaction of Cel7A with cellulose and the velocity and run length of molecules that have already entered a processive state, further emphasizing that adsorption, productive complexation, processive hydrolysis, stalling, and dissociation represent mechanistically distinguishable steps ^16^. The present SANS measurements do not distinguish individual productive and nonproductive molecules, and a heterogeneous population is expected even under conditions that support catalysis. Nevertheless, the emergence of a distinct interdomain correlation specifically under catalytically favorable conditions suggests that productive engagement with cellulose is associated with a redistribution of the conformational ensemble of the intact holoenzyme. We therefore interpret the active cellulose-bound state not as a single structure, but as a substrate-selected conformational ensemble that is enriched under conditions favorable for catalysis.

The SANS data further indicate that this substrate-selected ensemble involves the relative organization of the CD, linker, and CBM rather than a simple global collapse of the protein. The principal peak in the *p(R)* distribution at approximately 24 Å is present under all conditions and is largely attributable to intradomain correlations within the catalytic domain, whereas the longer-distance features report on correlations involving the linker and CBM. At pD 4.2 and 5.3, the appearance of the additional correlation at approximately 46 Å is consistent with an increased population in which part of the linker/CBM region occupies a more defined position relative to the catalytic domain. At the same time, the longer-distance component of the *p(R)* distribution remains evident, indicating that extended conformations continue to be populated. Earlier molecular dynamics simulations of an isolated Cel7A linker identified free-energy minima corresponding to compressed and extended linker configurations of approximately 25 and 55 Å, respectively ^19^. The approximately 22–24 Å separation between the principal and secondary peaks observed here under catalytically favorable conditions is therefore comparable to the length predicted for a compressed linker configuration. However, rather than implying interconversion between two rigid conformations, the present data are better interpreted as a shift in the relative populations of a heterogeneous conformational ensemble.

This ensemble-based interpretation is also consistent with recent work on the Cel7A linker and its extensive glycosylation. The linker has traditionally been viewed primarily as a flexible tether connecting the CD and CBM, and deletion or removal of the linker reduces both cellulose binding and catalytic activity ^20^. However, the glycosylated linker itself can interact dynamically with cellulose. Payne et al. showed that addition of the native glycosylated Cel7A linker to CBM1 increased its affinity for crystalline cellulose by approximately an order of magnitude compared with the isolated CBM, while molecular simulations indicated transient association of linker glycans with the cellulose surface ^58^. Integrative modeling and molecular dynamics of full-length Cel7A subsequently demonstrated that O-glycosylation restricts the conformational ensemble of the linker, favors more elongated structures, and results in more coordinated interdomain motion. Whereas in the absence of glycosylation, substantially greater conformational plasticity and frequent compression of the CD and CBM were observed ^21^. These studies support the view that the linker is not simply a passive tether or mechanical spring. Instead, linker flexibility and glycosylation appear to define the range and probability of interdomain configurations available to the holoenzyme and may thereby influence its interaction with the cellulose interface. The substrate-dependent changes observed in the present SANS measurements are consistent with this more dynamic role and suggest that the cellulose surface selectively stabilizes or enriches particular interdomain arrangements from the broader conformational landscape of Cel7A.

The results also provide an opportunity to reconsider the classical “caterpillar” model proposed for Cel7A processivity. In the original model, both the CD and CBM remain associated with the same cellulose chain and coordinated compression and extension of the linker allow the domains to advance alternately along the substrate ^17–19^. Such a model is attractive because it provides a mechanical role for the multidomain architecture of the enzyme. However, subsequent experiments have demonstrated that the essential processive movement is intrinsic to the catalytic domain. High-speed atomic force microscopy directly visualized Cel7A moving unidirectionally along crystalline cellulose at approximately 3.5 nm s⁻¹ and showed that removal of the CBM did not eliminate processive movement, indicating that the catalytic domain contains the principal machinery required for chain engagement and translocation ^14^. Current models of processive glycoside hydrolases likewise emphasize the energetic coupling among polymer-chain complexation, decrystallization, movement through the catalytic tunnel, hydrolysis, product release, and decomplexation rather than requiring a mechanically stepping CBM ^1^.

Our SANS-constrained simulations are consistent with this revised view. None of the conformations sampled in simulations of Cel7A free in solution accurately reproduced the secondary correlation observed experimentally for cellulose-bound Cel7A under active conditions, suggesting that interaction with the cellulose interface stabilizes or enriches conformations that are poorly sampled in solution. Furthermore, simulations in which the catalytic cleft faced the cellulose surface and the CD and CBM were constrained to the same cellulose strand, while their separation was varied over a broad range, did not reproduce the secondary feature of the experimental *p(R)* distributions. Thus, the data are not readily explained by a simple configuration in which the CD and CBM remain simultaneously constrained to a single cellulose chain throughout the catalytic cycle. Importantly, this result does not exclude coordinated motions of the two domains or transient simultaneous surface contact, nor does it establish a unique alternative geometry. Rather, it argues against the most restrictive interpretation of the caterpillar mechanism in which the CBM that mechanically advances with the catalytic domain along the same cellulose strand.

A more flexible model is consistent with both the present results and the recent literature. The CBM may primarily increase the probability that Cel7A remains localized at the cellulose interface, while the glycosylated linker permits the catalytic domain to explore different positions and orientations relative to the surface. This arrangement would allow the CD to engage accessible reducing chain ends or chain breaks while the CBM remains associated elsewhere on the fibril. Earlier biochemical studies showing that the linker and CBM increase cellulose affinity and affect catalytic efficiency are compatible with such a role ^20,58^. At the single-molecule level, the existence of static, processive, and stalled surface-bound populations further suggests that productive engagement is not determined simply by whether Cel7A is adsorbed to cellulose, but by whether the catalytic domain can adopt a geometry that permits chain complexation and entry into the catalytic tunnel ^15,16^. The enhanced interdomain correlation observed here may therefore reflect a substrate-selected configuration, or family of configurations, associated with this transition. Some of the MD conformations consistent with the additional *p(R)* feature contain a pronounced bend in the linker, raising the possibility that the CD can transiently move away from direct contact with the cellulose surface while the CBM remains associated. Our data do not require such detachment, however, the low spatial resolution and ensemble nature of SANS preclude assignment of a unique trajectory. We therefore regard transient CD repositioning or detachment as a mechanistic model consistent with the observations rather than as a direct structural conclusion.

The substrate-dependent conformational selection observed here may reflect a more general requirement for enzymes acting on recalcitrant polymers to reorganize upon productive engagement with a solid interface. Processive chitinases provide a particularly close mechanistic comparison. Like Cel7A, *Serratia marcescens* chitinase A acts processively on a crystalline polysaccharide using an extended substrate-binding cleft. Single-molecule measurements showed approximately 1-nm forward and backward steps during processive movement and demonstrated that extraction, or decrystallization, of a single chitin chain from the crystal surface is considerably slower than the glycosidic-bond cleavage reaction itself ^2^. Thus, for both cellulose and chitin degradation, productive catalysis requires coupling active-site chemistry to the physical problems of polymer-chain capture, decrystallization, translocation, and maintenance of a catalytically competent interfacial complex. In this context, the structural redistribution observed for Cel7A may represent one manifestation of a broader mechanistic principle: interaction with a heterogeneous polymer surface selects among multiple enzyme conformations, only a subset of which are compatible with productive substrate engagement.

A similar distinction between adsorption and productive engagement is emerging for enzymes that degrade synthetic insoluble polymers. Fluorescence measurements of PET hydrolases have revealed slow adsorption/desorption dynamics and long-lived surface-associated populations, highlighting that strong or persistent surface association is not necessarily equivalent to productive catalysis ^59^. Although PET hydrolases are not generally considered processive like Cel7A, they face the same requirement to transition from nonspecific or weakly productive surface association into a geometry that permits polymer-chain engagement and bond cleavage. These systems therefore reinforce the broader relevance of distinguishing among solution, adsorbed, productively engaged, stalled or nonproductive, and dissociated states when considering enzymes that act at insoluble polymer interfaces.

Several limitations should be considered when interpreting the present results. First, SANS measures a population-averaged conformational ensemble and cannot identify the catalytic state of individual surface-bound molecules. The samples therefore almost certainly contain mixtures of adsorbed, productively engaged, processive, stalled, and nonproductive Cel7A, even at pD values that favor cellulose hydrolysis. Second, alteration of pD changes the conformational equilibrium of Cel7A in solution as well as its catalytic activity, making it difficult to assign every structural difference exclusively to catalysis. Third, the structural information obtained from SANS is low resolution, so individual domain orientations and transient contacts cannot be uniquely assigned. Finally, the constrained MD simulations test defined classes of surface-bound configurations, so failure of a particular configuration to reproduce the experimental scattering excludes that model under the imposed constraints but does not uniquely specify the geometry of the productive complex. These limitations argue for an ensemble-level interpretation of the data and against assigning the scattering observations to a single molecular trajectory.

## Conclusions

The combined contrast-variation SANS and molecular dynamics results show that productive engagement of *T. reesei* Cel7A with cellulose is accompanied by its reorganization into a substrate-selected ensemble that is distinct from the conformations sampled in solution. Under catalytically favorable conditions, cellulose-bound Cel7A exhibits structural correlations not observed for the free enzyme or for the poorly active bound state, indicating that adsorption alone is insufficient to define the catalytically relevant configuration. The inability of models in which the catalytic domain and CBM are constrained to the same cellulose strand, to reproduce the bound-state scattering argues against a simple rigid caterpillar-like mechanism and instead supports a more dynamic model in which the CBM maintains interfacial localization while the glycosylated linker allows the catalytic domain to explore positions and orientations compatible with productive chain engagement, processive hydrolysis, and release. More broadly, these findings support the view that heterogeneous biocatalysis is governed not only by active-site chemistry and substrate affinity, but also by interface-selected conformational states that determine whether binding becomes productive. The study also demonstrates the advantage of neutron scattering using biosynthetic deuteration and contrast matching. Nanoscale structural information can be obtained for proteins operating at complex interfaces under functionally relevant conditions. This approach should be broadly applicable to other enzymes and biomolecules that function at insoluble polymer surfaces and in heterogeneous biological environments, where the conformational states most relevant to activity may be inaccessible to conventional structural methods.

## Supporting Information

Additional details on determination of the contrast match point for deuterated cellulose; binding isotherms of Cel7A on bacterial cellulose; temperature dependent hydrolysis of bacterial cellulose by Cel7A; SANS analysis of cellulose-cellulase interactions; and molecular dynamics simulations

## Author contributions

SVP performed and analyzed SANS experiments with input from WTH, VSU, and PL; JH, BRE, BHD and HO’N designed and performed experimental studies; LP, EA, XC, BL and JCS designed and performed computational studies; SVP and HO’N wrote the manuscript with input from all authors. All authors had the opportunity to read and comment on the manuscript.

## Acknowledgements

This research is funded by the Genomic Science Program, Office of Biological and Environmental Research, U. S. Department of Energy, under FWP ERKP752. The Center for Structural Molecular Biology operates Bio-SANS and is supported by the U.S. DOE, Office of Science, Office of Biological and Environmental Research under FWP ERKP291. A portion of this research used resources at the High Flux Isotope Reactor and Spallation Neutron Source, DOE Office of Science User Facilities operated by the Oak Ridge National Laboratory. The beam time was allocated to Bio-SANS on proposal number IPTS-IPTS-6649.1. An award of computer time was provided by the INCITE program. This research used resources of the OCLF, which is supported under contract DE-AC05-00OR22725. Oak Ridge National Laboratory (ORNL) is managed by UT-Battelle, LLC, for the U. S. Department of Energy (DOE) under contract No. DE-AC05-00OR22725.

---

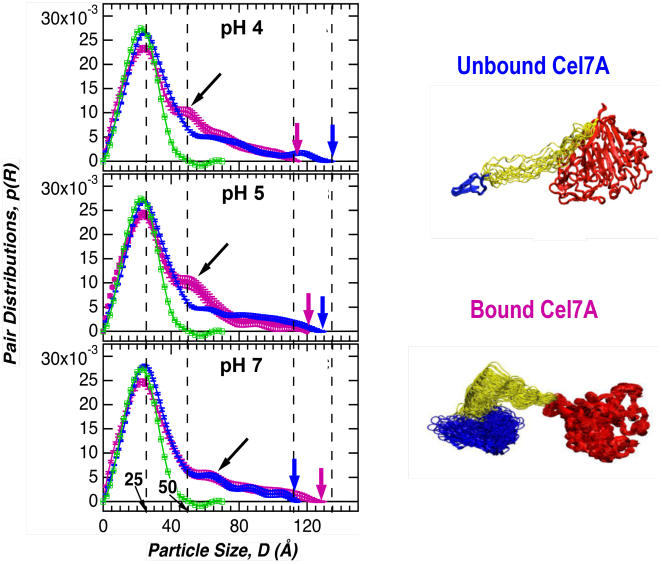

