## Supporting Information for "Conformational Reorganization of a Processive Cellulase on Crystalline Cellulose"

**Determination of the contrast match point for deuterated cellulose:** Deuterated cellulose produced as described previously [1,2], was equilibrated in different $D_2O/H_2O$ ratios (0–100%) and the SANS scattering profiles were measured (Figure S1A). A plot of $\sqrt{I(Q)}$ versus % $D_2O$ at $Q$ = 0.004 Å$^{-1}$ after background subtraction is shown in Figure S1B. The contrast match point was determined to be 85.4 ± 1.5 % $D_2O$. The experimentally determined contrast match point was largely $Q$ independent over the measured $Q$ range evident from all the curves being parallel to each other over the entire $Q$-range as shown in Figure S1A. Also, the scattering profile of the cellulose measured at the contrast match point was strongly attenuated over the measured $Q$-range compared to the material in 100% $D_2O$. The monotonously increasing scattering intensity with decreasing wave-vector $Q$ observed in the low-$Q$ region may be due to the presence of amorphous and crystalline regions in the cellulose or the presence of regions of the cellulose matrix that are inaccessible to the solvent.

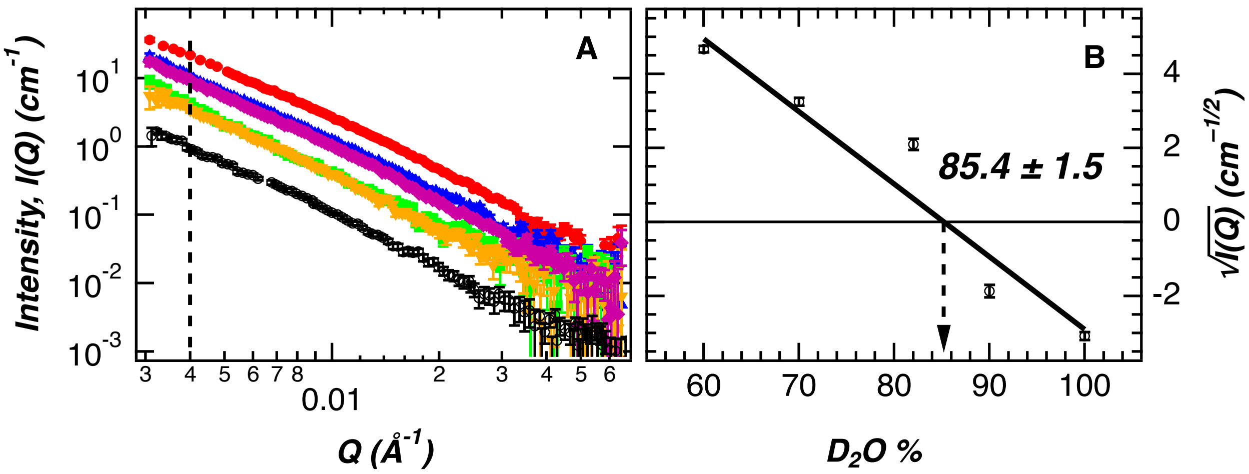


**Figure S1.** (A) Contrast variation SANS profiles of deuterated bacterial cellulose. Solvent ratios in % $D_2O$ used for the measurements were 60 (red), 70 (blue), 82 (green), 90 (orange) and 100 (purple). Scattering data in contrast matching buffer, 85.4 % $D_2O$ (black) is also included and the dashed line shows Q = 0.004 Å$^{-1}$. (B) Determination of the contrast matching solvent ratio ($D_2O/H_2O$) from the trend of the scattering intensity square root (open black circles) at $Q$ = 0.004 Å$^{-1}$ with solvent ratio and black solid line is a linear fit. The contrast matching point was determined to be 85.4 ± 1.5 % $D_2O$.

**Adsorption of Cel7A on bacterial cellulose:** The optimal time for binding of *T. reesei* Cel7A to the bacterial cellulose was 30 min as determined by mixing 5 µmol (0.335 mg/mL) of Cel7A with 0.1% (final conc.) bacterial cellulose and incubating the reaction at 15, 20 and 30 °C for time periods ranging from 15 to 180 min. The samples were treated as described below. To calculate a binding isotherm for the interaction of *T. reesei* Cel7A with bacterial cellulose, samples of bacterial cellulose (0.1% w/v) were incubated with different concentrations ($A_{280nm}$ = 0.02 to 1.17) of *T. reesei* Cel7A in 0.2 mL of 50 mM Na acetate, pD 5.3 using a shaking water bath (Lindberg/Blue M SWB1122A-1) for constant agitation. Each reaction condition was carried out in triplicate at 15, 20, and 30 °C. After 30 min, the samples were centrifuged at 16,000 x g for 60s to terminate the reaction, and the supernatants (100 µl) were collected immediately. The $A_{280nm}$ of the supernatant was measured and the free enzyme concentration was determined using

$E_{280nm}^{1M}$ =73,000 $M^{-1}cm^{-1}$, the previously reported extinction coefficient for *T. reesei* Cel7A [3].The amount of enzyme bound to cellulose was determined by subtraction of the amount of free enzyme from the total amount of enzyme divided by cellulose mass [4–8]. The apparent binding capacity and dissociation constant for the interaction of *T. reesei* Cel7A with bacterial cellulose were calculated from the fit to a plot of the amount of adsorbed enzyme (nmol/mg-cellulose) versus free enzyme concentration using the following equation [7]:

$$B = \frac{B_{max} \times F}{K_d + F} \quad (1)$$

Where $B$ is the amount of bound enzyme (nmol/mg-cellulose), $F$ is the amount of free enzyme (nM), $B_{max}$ is the maximum amount of adsorbed enzyme (nmol/mg-cellulose), and $K_d$ is the dissociation constant. The amount of reducing sugar released during cellulose hydrolysis was measured with the 3,5-dinitrosalicylic acid (DNS) assay [4].

Figure S2A shows the curve for binding of Cel7A to cellulose and Figure S2B shows the Scatchard plot of the data. At approximately, 10 and 14 µM Cel7A the bacterial cellulose became saturated with the enzyme at 15 and 20 °C, respectively. In the case of the 30 °C data, less binding was observed due to increased enzymatic activity at the higher temperature resulting in rapid hydrolysis of the cellulose to soluble sugars. The Scatchard plots are linear for all three measured temperatures which indicates that the binding is of a Langmuir type. This allowed a one-site binding model to be used to fit the data. The adsorption parameters obtained using Equation 1 are summarized in Table S1. The maximum amount of bound enzyme was 7.35 nmol/mg cellulose at 20 °C, which was 1.49 and 5.36 times higher than the values obtained at 15 and 30 °C, respectively. Both the one-site binding model and Scatchard analysis yielded similar values for the maximum amount of adsorbed enzyme at each temperature

**Table S1:** The temperature dependent binding capacities and the association and dissociation constants for the adsorption of *T.reesei* Cel7A to bacterial cellulose

| Temp (°C) | One-Site Binding Model | | Scatchard Analysis | |
|---|---|---|---|---|
| | Dissociation constant (µM) | Binding capacities (nmol Cel7A / mg cellulose) | Association constant ($\mu M^{-1}$) | Binding capacities (nmol Cel7A / mg cellulose) |
| 15 | 1.48±0.11 | 4.91±0.13 | 0.60±0.05 | 5.27±0.08 |
| 20 | 1.94±0.10 | 7.35±0.13 | 0.58±0.04 | 7.08±0.08 |
| 30 | 0.23±0.09 | 1.37±0.13 | 1.57±0.20 | 2.37±0.20 |

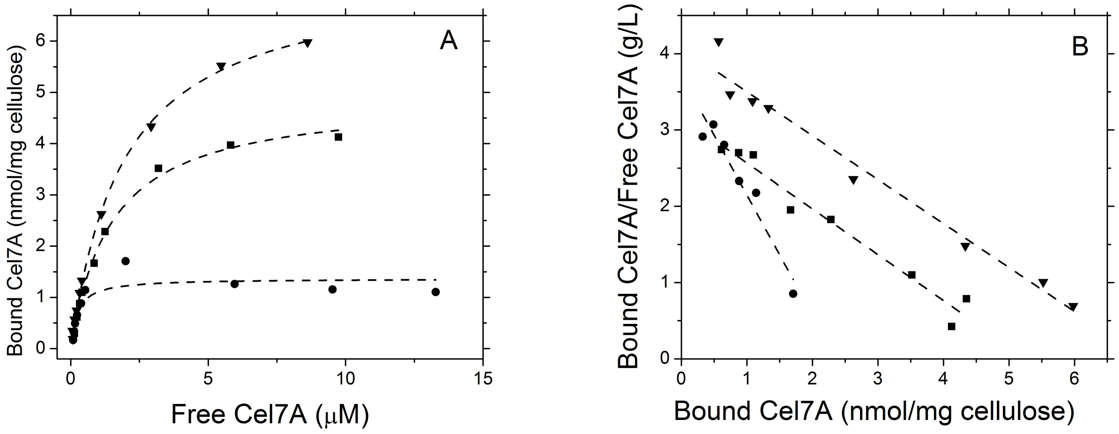


**Figure S2.** The concentration-dependent binding of *T. reesei* Cel7A to bacterial cellulose as a function of temperature. *Left panel:* The temperature dependence of the adsorption isotherms of Cel7A binding to cellulose. *Right panel:* Scatchard analysis of the experimental data. 10 mg/mL bacterial cellulose was incubated with successively higher concentrations of protein in 50 mM Na acetate buffer pH 5.0, at 15(■), 20 (▼) and 30 °C (●). After 30 min the reaction was centrifuged and the amount of protein in the supernatant was determined. The experimental points are the mean values of triplicate determinations. The lines show the best fits obtained using a one-site binding model (*left panel*) and linear regression analysis (*right panel*)

**Temperature dependent hydrolysis of bacterial cellulose by Cel7A:** The concentration dependent hydrolytic rate of the cellulose by Cel7A was determined for each temperature after a 15 min incubation (Figure S3). The data indicate that the Cel7A hydrolyzed 11.4, 30.1 and 77% of the cellulose at 15, 20 and 30 °C respectively, at the highest concentration of Cel7A studied (14 μM, 1 mg/mL, 0.1 % cellulose, pD 5.3)

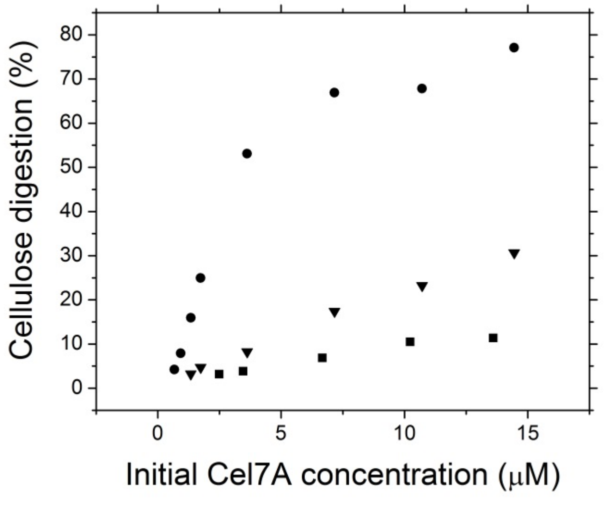


**Figure S3.** The effect of temperature on the bacterial cellulose hydrolysis at different concentrations of Cel7A. The protein was incubated with the cellulose in 50 mM Na acetate buffer pD 5.3 for 15 min at 15(■), 20 (▼) and 30 °C (●). The amount of reducing sugar that was released was measured using the DNS assay.

***SANS analysis of cellulose-cellulase interactions:*** The radius of gyration, $R_g$, was determined according to Guinier and Fournet [9] where the scattering intensity and wave vector $Q$ are related by the expression in Equation 2,

$$ln[I(Q)] = \left(-\frac{R_g^2}{3}\right) Q^2 + ln[I_0] \quad (2)$$

The slope of a simple linear fit ($y = mx + c$) to the curve in the plot of $ln[I(Q)]$ vs. $Q^2$ provides an estimate of the $R_g$ that is valid only if the small-angle scattering condition is satisfied, $Q_{max} \times R_g < 1.3$. The indirect Fourier transform was used to determine the pair-correlation function $p(r)$ implemented in the software package GNOM [10] using the measured intensity profiles (Table S2). The boundary conditions imposed were $p(r) = 0$ at r = 0 and $D_{max}$ (maximum linear dimension). Structural modeling of Cel7A from the SANS data was accomplished using DAMMIN [11]. All data were modeled over the same $Q$-range to provide consistency in the information content in the data. A set of 37 models was generated using default settings from each SANS dataset and a consensus envelope, representing the structure consistent across the set of models was determined using DAMAVER program suite [12].

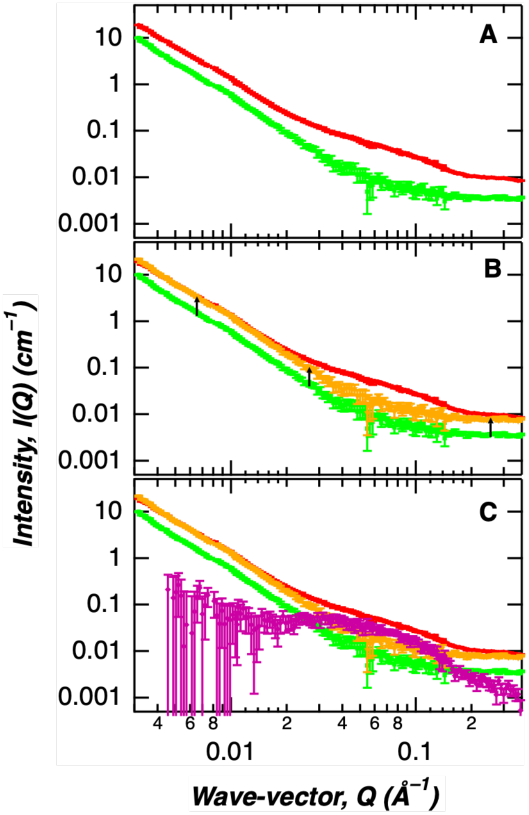


**Figure S4.** Steps used to extract Cel7A enzyme's small-angle neutron scattering profile when bound to bacterial cellulose substrate. (A) Scattering profiles of Cel7A-cellulose complex (red) and cellulose only (green). (B) Scaled scattering intensity of cellulose only curve (orange) to match low $Q$ scattering intensity which represents bacterial cellulose substrate in Cel7A-cellulose complex. (C) Scattering profile of only bound Cel7A (purple) after subtraction of residual scattering signal from bacterial cellulose (purple).

The partially deuterated bacterial cellulose scattering was not fully matched over the entire $Q$-range in the contrast match buffer that was determined by contrast variation study (Figure S1). Instead, some residual scattering signal remained in the low-$Q$ region that is attributed to density heterogeneity within the cellulose matrix. Figure S4 summarizes the approach utilized to obtain SANS profiles of the bound Cel7A enzyme.

**Table S2.** Guinier analysis and GNOM program output and input parameters (Expanded version of Table 1)

| pD | Guinier | | | | | $p(R)$ | | |
|---|---|---|---|---|---|---|---|---|
| | $R_g$ (Å) | $\pm\Delta R_g$ (Å) | $Q_{min}$ (Å$^{-1}$) | $Q_{max}$ (Å$^{-1}$) | $Q_{max}$*$R_g$ | $R_g$ (Å) | $\pm\Delta R_g$ (Å) | $D_{max}$ (Å) |
| *Free* | | | | | | | | |
| 4.2 | 30.3 | 0.7 | 0.0142 | 0.0370 | 1.12 | 32.0 | 0.8 | 135 |
| 5.3 | 30.9 | 1.0 | 0.0084 | 0.0324 | 1.00 | 32.6 | 0.7 | 130 |
| 7.0 | 29.3 | 0.8 | 0.0086 | 0.0358 | 1.05 | 29.4 | 0.6 | 115 |
| *Bound* | | | | | | | | |
| 4.2 | 32 | 9 | 0.0081 | 0.0515 | 1.00 | 30.5 | 1.3 | 115 |
| 5.3 | 33 | 8 | 0.0070 | 0.0335 | 1.09 | 30.2 | 2.2 | 120 |
| 7.0 | 33 | 9 | 0.0065 | 0.0314 | 1.04 | 31.4 | 1.7 | 130 |

**Molecular Dynamics Simulations:** The sequence of the linker peptide is PGPSSGTTTAPRRTTTTGPPNGGPPNG [13]. The linker modeled in the fully extended form has a length of approx. 92 Å. Three N-linked glycosylation sites of *T. reesei* Cel7A (N45, N270 and N384) have been identified experimentally [13]. Glycosylation patterns, which affect enzyme activity, vary considerably depending on the microbial growth conditions. In this MD study, the glycosylation of the linker was based on the suggestion of Harrison and coworkers [13]. The CD residues N270 and N384 were bonded to $Man_5GlcNAc_2$ while the CD residue N45 was not glycosylated.

Five MD simulations of Cel7A in explicit solvent were performed. 20 sodium and 3 chloride ions were added to neutralize each system and to create an ionic concentration of 100 mM. MD simulations were performed with NAMD [14] using the CHARMM27 force field for proteins [15–17], the C35 force field for carbohydrates [18,19], and the TIP3P model for water [20]. Periodic boundary conditions were used. The long-range electrostatic interactions were computed using the Particle Mesh Ewald method [21], for which the reciprocal sum was computed with sixth-order interpolation and 1.5 Å Fourier grid spacing. Van der Waals interactions were computed with a smooth switching function between 8 Å and the cutoff value of 10 Å.

The MD systems were first energy-minimized for 5000 steps using the conjugate gradient algorithm and then gradually heated up from 50 K to 300 K over 40 ps. During the minimization and heating phases, harmonic restraints with a force constant of 5 kcal/(mol·Å$^2$) were applied to the backbone atoms. The heated systems then underwent four equilibration steps. In the first 200 ps equilibration run, all heavy protein atoms were fixed to relax the solvent. In the remaining three equilibration runs, each 200 ps long, the harmonic restraints were gradually lifted from 5 to 1 kcal/(mol·Å$^2$). Langevin dynamics was used to maintain constant temperature at 300 K and the Nosé-Hoover Langevin piston [22] with a decay period of 500 fs was used for maintaining constant pressure at 1.01325 bar. After equilibration, production runs were performed for 60 to 250 ns at isobaric-isothermal (NPT) ensemble. A time step of 1 fs was used for numerical integration of the equation of motion. Coordinates were saved every 1 ps. A total of five MD trajectories with different initial configurations (velocities) were produced; the lengths for individual simulations are 464 ns, 375 ns, 377 ns, 465 ns and 319 ns, respectively, resulting in a total simulation time of approx. 2 microseconds.

**Clustering:** RMSD-based clustering was performed for a total of 20004 frames using *g_cluster* with gromos algorithm. Different RMSD cutoffs from 3 to 9 Å were tested, and the most sensible set of results (in terms of number of clusters and cluster populations) were found when the RMSD cutoff was set to 6 Å, which resulted in 36 initial clusters. As discussed below, the final number of clusters was reduced to 11 to ensure each cluster contains > 10 frames and has a distinct theoretical $I(Q)$ profile (Figure S5).

***I*(*Q*) calculation:** For each cluster, the scattering intensity $I(Q)$ was calculated using SASSENA [23]. To match the experimental contrast variation condition (85% $D_2O$ and 15% $H_2O$), 85% of both water and Cel7A exchangeable hydrogen atoms were randomly replaced by deuterium atoms. Solvent molecules were included in the SANS in the calculation.

**Minimal set of *I*(*Q*) curves:** To obtain a minimal set of $I(Q)$ curves, we removed clusters which contain <10 frames because they represent poor configurational statistics. The similarity between the $I(Q)$ curves of two different clusters were accessed by computing $\varepsilon_{ij}$ defined by the following equation

$$\varepsilon_{ij} = \sum_Q \left[I_i(Q) - I_j(Q)\right]^2 \qquad (3)$$

for all pairs of clusters *i* and *j*. For cases where $\varepsilon_{ij} < 0.06$, the clusters *i* and *j* were considered to have similar $I(Q)$ curves, and the cluster with lower number of frames was removed. This ensures that each of the final 11 clusters have a distinct $I(Q)$ curve (Figure S5).

**Scaling of Experimental and MD *I*(*Q*) Curves:** The absolute scales of the experimental $I_E(Q)$ and theoretical $I_T(Q)$ curves are different. Therefore, comparison between theoretical and the simulation data requires scaling the experimental $I(Q)$ data, $I_E^{Sc}(Q)$, using the following scaling scheme:

$$I_E^{Sc}(Q) = C + B \times I_E(Q) \qquad (4)$$

where constants $B$ and C were adjusted to ensure the experimental and theoretical data span the same intensity range. To find the optimal values for the scaling parameters $B$ and $C$, a Monte Carlo search was performed such that the difference between the experimental and simulated data was minimized. The optimization was done for all MD clusters and the average values of the scaling parameters, $B$ and $C$, were calculated for each of the six experimental $I(Q)$ curves, $I_E(Q)$ and presented in Table S5.

**Table S5.** Scaling parameter values to match the scattering intensity range for experimental and theoretical profiles using Equation 4.

| Samples | B | C |
|---|---|---|
| Free, pD 4.2 | 8.1749 | 0.005 |
| Free, pD 5.3 | 8.2762 | 0.005 |
| Free, pD 7.0 | 8.5074 | 0.002 |
| Bound, pD 4.2 | 15.0545 | 0.015 |
| Bound, pD 5.3 | 15.1105 | 0.015 |
| Bound, pD 7.0 | 14.9425 | 0.015 |

**Optimization:** To find the best fit to the experimental $I(Q)$, the theoretical SANS curve $I_T(Q)$ was modeled as a linear combination of the minimal set of eleven $I_n(Q)$ curves:

$$I_T(Q) = \sum_{n=1}^{11} p_n I_n(Q) \tag{5}$$

with the following two constraints on the to-be determined coefficients $a_i$:

$$0 \leq a_n \leq 1 \tag{6}$$

$$\sum_n a_n = 1 \tag{7}$$

The optimization aims at finding a set of $p_n$ values that minimize the following function:

$$\Delta I = |I_E^{Sc} - I_T| \tag{8}$$

The Levenberg–Marquardt algorithm (LMA) was used for optimization because it is well studied and very fast (1000 iterations takes <5 seconds on a 1.6 GHz laptop). Nevertheless, like most other optimization algorithms, LMA only finds local minima rather than the global minimum. The GNU R library implementation of the algorithm "minpack.lm" was used in the present study.

Several approaches are available to deal with the second constraint (Equation 7), and each implementation might have a different influence on the numerical stability of the LMA algorithm. To find the implementation with the best numerical stability, a four-step procedure was employed. In the first step, the optimization was performed using three different random initial populations $p_n$'s starting parameters without the constraint (Equation 7) to determine which cluster contributes the most to $I_T(Q)$. In the second step, the constraint (Equation 7) was enforced in a way that the largest population cluster, *m,* was chosen as "free parameter":

$$p_m = 1 - \sum_{n \neq m}^{11} p_n \tag{9}$$

As discussed below the choice of the free parameter does not influence the final populations. In the third step, clusters with a population <1% were removed in an iterative manner until the algorithm produced populations that were all > 1%. In the fourth step, to ensure that the results do not depend on the choice of "free parameter", steps 2 and 3 were repeated using another cluster as a free parameter than the one chosen in step 2. Importantly, the converged result was independent of the free parameter choice. The resulting $I(Q)$ plot is shown in Figure S6.

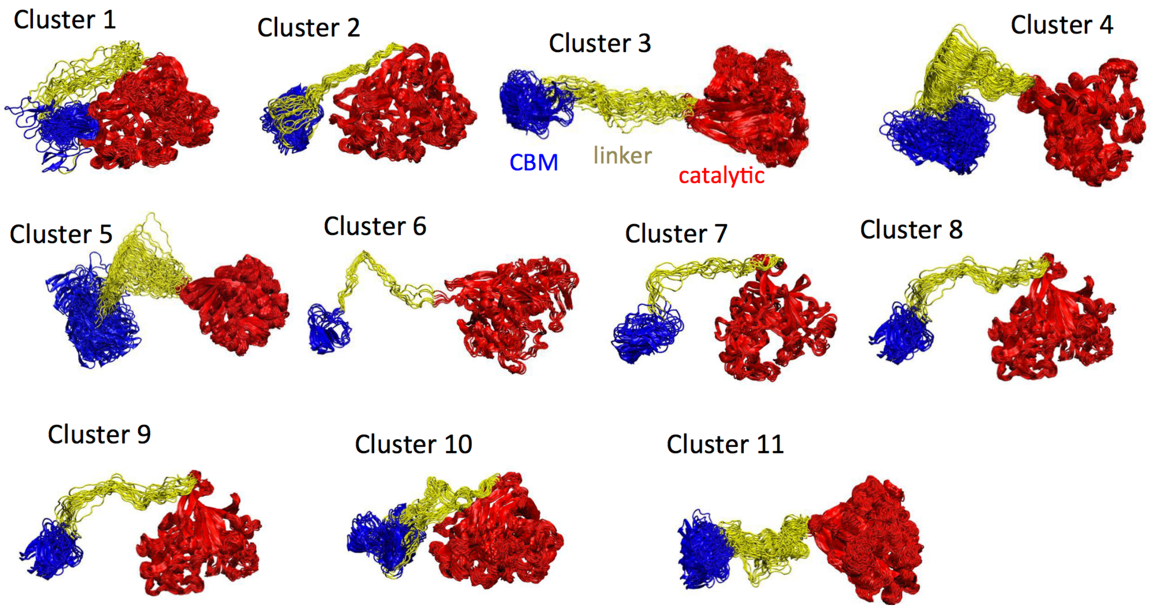


**Figure S5.** The 11 clusters sampled during the unconstrained MD simulations of free Cel7A in solution.

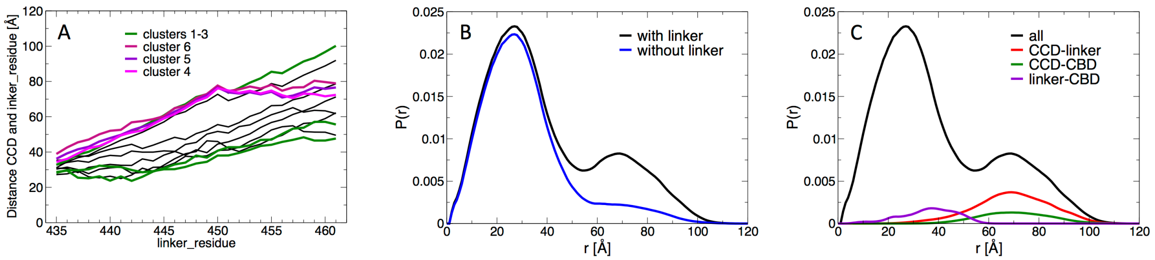


**Figure S6.** (A) Distance between the center of mass of the CCD domain and a linker residue, plotted for all 11 clusters in Figure S5. (B) $p(R)$ plot of cluster 4. “with linker” contains correlations between all atoms (CCD and linker and CBD), “no linker” excludes the linker atoms from the correlations (CCD and CBD). (C) $p(R)$ plot of cluster 4. “all” contains intra- and inter-domain contributions. CCD-linker, CCD-CBD and linker-CBD contain only the respective inter-domain contributions.

(17) MacKerell, A. D.; Bashford, D.; Bellott, M.; Dunbrack, R. L.; Evanseck, J. D.; Field, M. J.; Fischer, S.; Gao, J.; Guo, H.; Ha, S.; Joseph-McCarthy, D.; Kuchnir, L.; Kuczera, K.; Lau, F. T. K.; Mattos, C.; Michnick, S.; Ngo, T.; Nguyen, D. T.; Prodhom, B.; Reiher, W. E.; Roux, B.; Schlenkrich, M.; Smith, J. C.; Stote, R.; Straub, J.; Watanabe, M.; Wiórkiewicz-Kuczera, J.; Yin, D.; Karplus, M. All-Atom

Empirical Potential for Molecular Modeling and Dynamics Studies of Proteins. *J. Phys. Chem. B* **1998**, *102* (18), 3586–3616. https://doi.org/10.1021/jp973084f
(18) Guvench, O.; Mallajosyula, S. S.; Raman, E. P.; Hatcher, E.; Vanommeslaeghe, K.; Foster, T. J.; Jamison, F. W.; MacKerell, A. D. CHARMM Additive All-Atom Force Field for Carbohydrate Derivatives and Its Utility in Polysaccharide and Carbohydrate–Protein Modeling. *J. Chem. Theory Comput.* **2011**, *7* (10), 3162–3180. https://doi.org/10.1021/ct200328p
(19) Guvench, O.; Greene, S. N.; Kamath, G.; Brady, J. W.; Venable, R. M.; Pastor, R. W.; Mackerell, A. D. Additive Empirical Force Field for Hexopyranose Monosaccharides. *J. Comput. Chem.* **2008**, *29* (15), 2543–2564. https://doi.org/10.1002/jcc.21004
(20) Jorgensen, W. L.; Chandrasekhar, J.; Madura, J. D.; Impey, R. W.; Klein, M. L. Comparison of Simple Potential Functions for Simulating Liquid Water. *J. Chem. Phys.* **1983**, *79* (2), 926–935. https://doi.org/10.1063/1.445869
(21) Darden, T.; York, D.; Pedersen, L. Particle Mesh Ewald: An $N \cdot \log(N)$ Method for Ewald Sums in Large Systems. *J. Chem. Phys.* **1993**, *98* (12), 10089–10092. https://doi.org/10.1063/1.464397
(22) Feller, S. E.; Zhang, Y.; Pastor, R. W.; Brooks, B. R. Constant Pressure Molecular Dynamics Simulation: The Langevin Piston Method. *J. Chem. Phys.* **1995**, *103* (11), 4613–4621. https://doi.org/10.1063/1.470648
(23) Lindner, B.; Smith, J. C. Sassena — X-Ray and Neutron Scattering Calculated from Molecular Dynamics Trajectories Using Massively Parallel Computers. *Comput. Phys. Commun.* **2012**, *183* (7), 1491–1501. https://doi.org/10.1016/j.cpc.2012.02.010
